# An MM and QM Study of Biomimetic Catalysis of Diels-Adler Reactions Using Cyclodextrins

**DOI:** 10.1101/169565

**Authors:** Wei Chen, Lipeng Sun, Zhiye Tang, Chia-en A. Chang

## Abstract

We performed computational research to investigate the mechanism by which cyclodextrins (CDs) catalyze Diels-Alder reactions between 9-anthracenemethanol and N-cyclohexylmaleimide. Hydrogen bonds (Hbonds) between N-cyclohexylmaleimide and the hydroxyl groups of cyclodextrins were suggested to play an important role in the catalysis.However, our free energy calculations and molecular dynamics simulations showed that these Hbonds are not stable, and quantum mechanics calculations suggested that the reaction is not promoted by these Hbonds. The binding of 9-anthracenemethanol and N-cyclohexylmaleimide to cyclodextrins was the key to the catalysis. Cyclodextrins act as a container to hold the two reactants in the cavity, pre-organizes them for the reactions, and thus reduces the entropy penalty to the activation free energy. Dimethyl-β-CD was a better catalyst for this specific reaction than β-CD because of its stronger van der Waals interaction with the pre-organized reactants and better performance in reducing the activation energy. This computational work sheds light on the mechanism of the catalytic reaction by cyclodextrins and introduces new perspectives of supramolecular catalysis.

## 1. Introduction

Supramolecular catalysts offer potential advantages over enzymes, including greater physical and chemical stability, lower molecular weight, and a far more varied selection of chemistries for the creation of structure and functionality. Due to their special chemical and physical properties, cyclodextrins are very useful in pharmaceuticals and food and agricultural industries [1-5]. Their enzyme-like hydrophobic pocket also allows it to catalyze chemical reactions just like an enzyme [6-12]. As a biomimetic supramolecular catalyst, β-cyclodextrins is also able to catalyze the important Diels-Alder reaction and synthesize various compounds [13-15].

The Diels-Alder reaction is an important carbon–carbon bond formation reaction in organic synthesis. It forms two carbon–carbon bonds and up to four new stereo centers in one step. For typical Diels-Alder reactions between diene and dienophile, frontier molecular orbital theory states that the interaction of the highest occupied molecular orbital (HOMO) of the diene with the lowest unoccupied molecular orbital (LUMO) of the dienophile is the dominant interaction in the transition state [16]. The rate of the Diels-Alder reaction could be accelerated by narrowing the energy gap between the HOMO and LUMO [17, 18].

Recently, cyclodextrins were reported to promote Diels-Alder reactions of 9- anthracenemethanol with a variety of N-substituted maleimides under mild reaction conditions [15]. In this paper, Chaudhuri *et al.* proposed a mechanism whereby the cyclodextrins bind the hydrophobic substituents on the maleimides (2a – 2d) and activate the dienophile by electronic modulation of the maleimide double bond via hydrogen bonds (Hbonds) with cyclodextrins (Figure 1). Methyl-β-cyclodextrin was found to bind maleimides with large N-substituents, such as 2a. more strongly than β-cyclodextrin, as indicated by the greater changes in the 1H NMR chemical shifts. Methyl-β-cyclodextrin is also significantly more efficient than β-cyclodextrin at promoting the reaction of N-cyclohexylmaleimide. The authors further suggested that this situation is likely because methyl-β-cyclodextrin is both more flexible and has more non-polar cavity than β-cyclodextrin [15]. However, if the reaction is promoted by the Hbonds between the maleimides and cyclodextrins, then β-cyclodextrin should perform at least as well as methyl-β-cyclodextrin because it has more hydroxyl groups to participate in the hydrogen bonding. How the flexibility and hydrophobicity of methyl-β-cyclodextrin helps the catalysis is unknown also.

**Figure 1.**
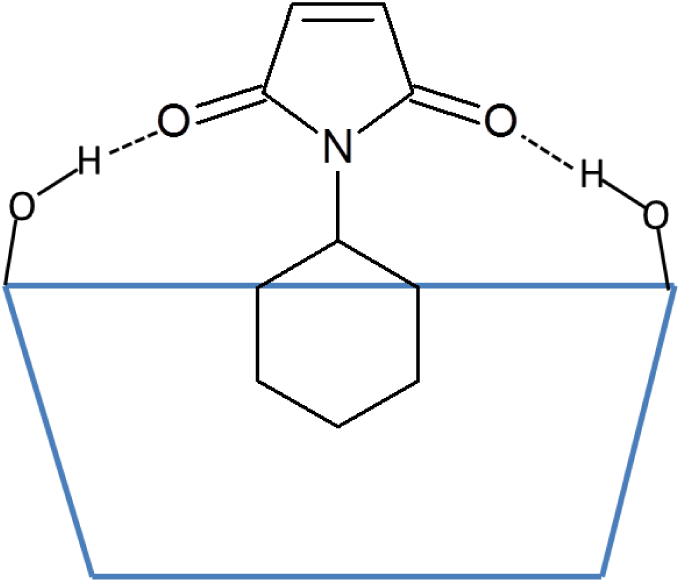
Illustration of the hydrogen bonds between the carbonyl groups on maleimides and the secondary hydroxyl groups on cyclodextrins. N-cyclohexylmaleimide is used as an example. The blue trapezoid represents cyclodextrins, and the dotted lines represent the hydrogen bonds.

In this paper, we performed computational work using both molecular mechanics, including free energy calculations and molecular dynamics (MD) simulations, and quantum mechanics tools to investigate Diels-Alder reactions with cyclodextrins as catalysts. We studied the reaction of 9- anthracenemethanol (compound 1) with N-cyclohexylmaleimide (compound 2a) (Figure 2) catalyzed by β-cyclodextrin. The study describes a new mechanism and explains why methyl-β-cyclodextrin outperforms β-cyclodextrin to promote the reactions.

**Figure 2.**
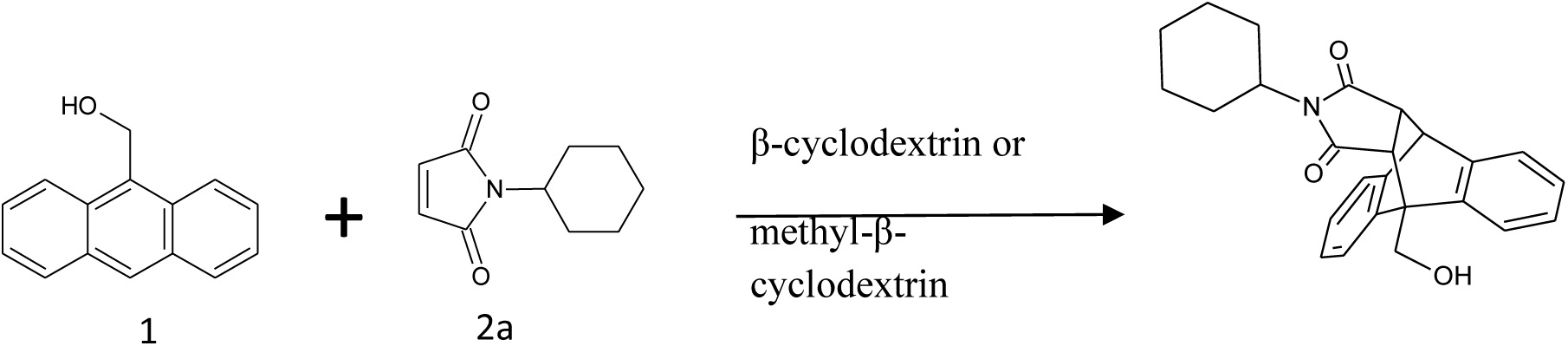
The Diels-Alder reaction of 9-anthracenemethanol (1) with N-cyclohexyl maleimide (2a) with β-cyclodextrin or methyl-β-cyclodextrin as catalyst.

## 2. Methods

Each α-D-glucopyranoside on cyclodextrins has one primary hydroxyl group and 2 secondary hydroxyl groups (Figure 3). All the commercially available methylated β-cyclodextrins are mixtures of various isomers and homologues except trimethyl β-cyclodextrin. Since the study by Chaudhuri *et al.*, showing an average of ∼1.8 methyl groups in methyl-β-cyclodextrin and the primary hydroxyl group more active than secondary hydroxyl groups in methylation, we assume that the major component of methyl-β-cyclodextrin is 2,6-O-dimethyl-β-cyclodextrin. For these reasons, we investigated the interactions of the two reactants (1 and 2a) in the Diels-Alder reaction with the catalysts β-cyclodextrin (β-CD) and 2,6-O-dimethyl-β-cyclodextrin (dimethyl-β-CD) by the computational methods briefly described below. For molecular mechanics calculations, force field q4md [19] and General Amber Force Field (GAFF) [20] were applied to cyclodextrins and the two reactants, respectively. The figures of molecule conformations in this paper were generated by using VMD [21].

**Figure 3.**
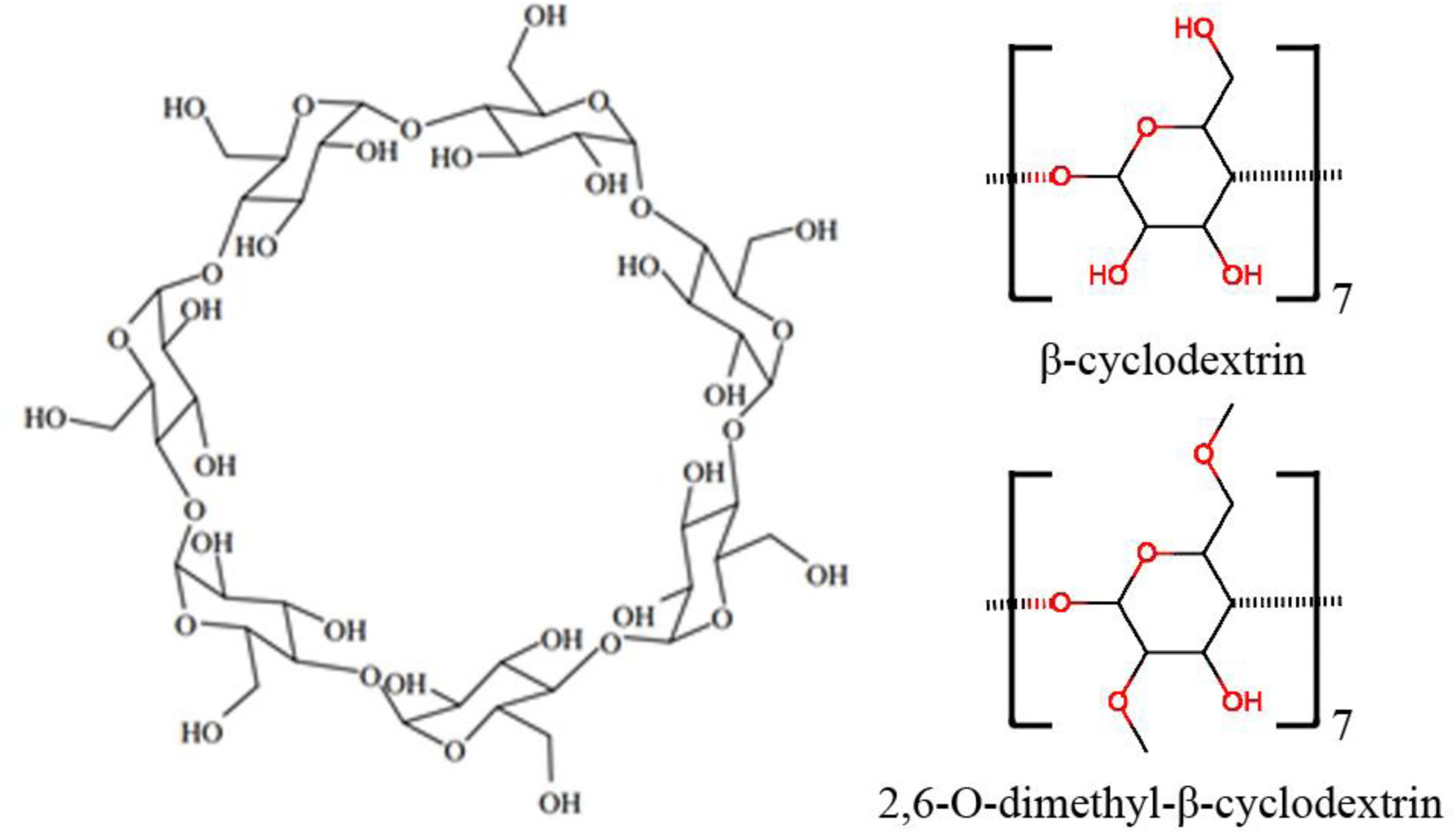
Molecular structure of β-cyclodextrin. The shape of the cyclodextrin macrocycle can be described as a truncated cone, with a narrow rim presenting primary hydroxyl groups and a wide rim presenting secondary hydroxyl groups on the glucose residues.

### 2.1 Free energy calculations with the VM2 method

The initial conformations of the complexes for the VM2 calculations were obtained by using an in-house docking script. The VM2 method [22] provides binding affinities by computing the standard chemical potentials of the free receptor, the free ligand, and their complexes and taking the differences to obtain the standard free energy of binding. The standard chemical potential of each molecular species is obtained as a sum of the contributions from the low-energy conformations of the species. These conformations are identified with the Tork conformational search algorithm [23], and a symmetry-corrected method ensures that no conformation is double-counted in the free energy sums [24]. The contribution of each unique energy well to the free energy is computed with an augmented form of the harmonic approximation, the Harmonic Approximation/Mode Scanning (HA/MS) method [25]. The ligand, the receptor, and their complex are treated as fully flexible during these calculations.

### 2.2 Unbiased MD simulations

We performed four sets of MD simulations, for 2a-β-CD complex, 2a-dimethyl-β-CD complex and pre-organized 1-2a reactant complexes binding to β-CD and dimethyl-β-CD, respectively. The initial conformations of the complexes of cyclodextrins and compound 2a for the MD simulations were the lowest energy conformations obtained from VM2. For the complexes of cyclodextrins with both reactants, the initial conformations were obtained by the following method: the reaction product was added in place of 2a in the initial conformations of the complexes of cyclodextrins and 2a, then the two single bonds connecting 1 and 2a in the product were removed and the whole system was energy minimized.

The Amber 14 package with GPU implementation [26, 27] was used for the MD simulations for the complexes. Minimization on the hydrogen atoms and the entire complex was applied for 500 and 5000 steps, respectively. After being solvated with a rectangular TIP3P water box [28], the edge of the box was at least 12 Å away from the solutes. The system went through a 1000-step water and 5000-step system minimization to correct any inconsistencies. Next, we relaxed the system by slowly heating it during an equilibrium course of 10 ps at 200, 250 and 298 K. We performed a production run in an isothermic-isobaric (NPT) ensemble with a 2-fs time step. The Langevin thermostat [29, 30], with a damping constant of 2 ps^-1^, was used to maintain a temperature of 298 K. The long-range electrostatic interactions were computed by the particle mesh Ewald method [31] beyond 8 Å distance. We collected the resulting trajectories every 2 ps. Finally, the SHAKE algorithm [32] was used to constrain water hydrogen atoms during the MD simulations. We performed 300 ns of MD production runs on each complex by using CPU parallel processing and local GPU machines. Finally, the trajectories were collected and analyzed at intervals of 20 ps.

In this work, an Hbond (X-H…Y) was considered formed if the distance between H and Y was smaller than 2.0 Å and the complimentary angle of X-H…Y was smaller than 90°(Figure SI-1). We used an in-house script to post-process the trajectories for direct Hbonds between 2a and cyclodextrins. The occurrence percentage of a Hbond was calculated as the number of the frames containing the Hbond divided by the total 3000 frames.

We computed the RMSD of compound 1 with respect to the initial conformation in the MD runs to determine whether compound 1 bound to the cavity of a β-CD or not. The RMSD data for compound 1 formed several clusters in this figure. The one with RMSD < 5 Å corresponded to the binding of 1 with 2a and β-CD/dimethyl-β-CD.

### 2.3 Quantum mechanics calculations with the pm3 method

Reaction paths and transition states (TSs) with and without β-CD and dimethyl-β-CD catalysts were calculated with the PM3 method in G09W package [33, 34]. Aqueous solvent was not included in the present calculations. The Gibbs free energies as well as the enthalpy and entropy components of reactants and TS were calculated with standard statistical thermodynamics equations [35] at 298K using the optimized molecular structures. For example, entropy was calculated with Eq. 1, where the partition function, *q*, is obtained by including contributions from translational, rotational, vibrational and electronic degrees of freedom.

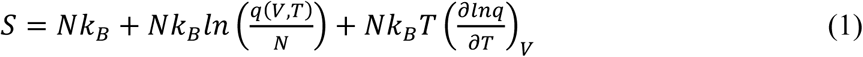

The activation free energies were obtained by computing the difference between the free energies of the reactants and the TS. Molecular structures were optimized by GEDIIS method [36] in G09.

We performed two sets of QM calculations for compounds 1 and 2a with β-CD or dimethyl-β-CD complexes with two β-CD conformations. The first set involved a conformation with Hbonds between the two carbonyl groups of 2a and the catalyst, and the other set did not have Hbonds between 2a and β-CD or dimethyl-β-CD. We also performed calculations for compounds 1 and 2a without β-CDs. Complex structures with the lowest energies in the MD runs were selected as initial structures for searching the TS of a cyclodextrin-catalyzed reaction and geometry optimization. The Berny algorithm [37, 38] was used for locating the TSs. After TSs were identified, the energy profile along the reaction coordinate connecting reactants, TS and products was calculated by the Intrinsic Reaction Coordinate (IRC). The step size was 0.02 amu^1/2^-Bohr and 150 steps were run on both forward and reverse directions along the reaction path.

## 3. Results and Discussion

### 3.1 Free energy calculation for cyclodextrin and substrate binding

To perform as a catalyst, cyclodextrin must form a close contact with a reactant(s). The cavity size of β-CD or dimethyl-β-CD is only large enough to accommodate one reactant, so we first searched for the reactant and β-CD–dimethyl-β-CD complex conformations and examined which reactant bound to a cyclodextrin first by computing the binding free energies of 1 or 2a to β-CD or dimethyl-β-CD with the VM2 method [22].

The calculated binding free energies and decompositions suggest that as the initial step of the catalyzed reaction, compound 2a is more likely to bind to cyclodextrins than compound 1 (Table 1). The computed binding affinity of 1 to β-CD and dimethyl-β-CD was −2.71 and −4.09 kcal/mol, at least 2 kcal/mol weaker than 2a binding to the catalysts. Compound 2a had a similar computed binding affinity with both β-CD and dimethyl-β-CD, −6.58 and −6.17 kcal/mol, respectively. Experimental NMR proton chemical shifts suggested that the binding of 2a with dimethyl-β-CD yielded a change in chemical shift by 0.078 ppm, as compared with 0.047 ppm from 2a binding with β-CD [15]. The NMR measurement did not explicitly quantify the binding affinity between 2a and β-CDs, but the changes in chemical shift suggested that 2a prefers binding to dimethyl-β-CD. Although our free energy calculations yielded similar binding free energy for 2a binding to both CDs, calculations show stronger enthalpic attraction between 2a and dimethyl-β-CD.

**Table 1.** Calculated free energies and breakdowns by the VM2 method for the complexes of reactants and cyclodextrins. Unit in kcal/mol.

| | $\Delta G_{\text{calc}}$ | $\Delta E_{\text{Valence}}$ | $\Delta E_{\text{Coulomb}}$ | $\Delta E_{\text{PB}}$ | $\Delta E_{\text{NP}}$ | $\Delta E_{\text{VDW}}$ | $\Delta H$ | $-T\Delta S$ |
| --- | --- | --- | --- | --- | --- | --- | --- | --- |
| β-CD and 1 | -2.71 | 1.96 | -8.04 | 15.69 | -2.86 | -25.27 | -18.52 | 15.81 |
| dimethyl-β-CD and 1 | -4.09 | 3.01 | -8.50 | 14.84 | -2.91 | -26.06 | -19.62 | 15.53 |
| β-CD and 2a | -6.58 | 0.39 | -5.98 | 13.23 | -2.76 | -25.12 | -20.23 | 13.65 |
| dimethyl-β-CD and 2a | -6.17 | 2.57 | -11.25 | 15.28 | -2.76 | -25.22 | -21.39 | 15.21 |

The binding modes computed by VM2 showed no significant Hbonds between compound 2a and cyclodextrins, so the Hbond formation may not be the functional structure of the catalyst. As shown in Figure 4, the cyclohexyl group of 2a snugly sits right in the hydrophobic cavity of cyclodextrins and the maleimide moiety stays at the wide rim side (the side with secondary hydroxyl groups) of cyclodextrins, whereas the anthracene rings stick into the cavity and remain vertical to the macrocycles of cyclodextrins. The principal axes of the reactants are aligned well with those of cyclodextrins to make tight contacts with each other. The main driving force for the binding is van der Waals interactions, which is expected because of the non-polar cavity of cyclodextrins. The value is similar for all complexes as well. Surprisingly, we observed no Hbonds between 2a and cyclodextrins in all conformations. In the complex with 2a, the secondary hydroxyl groups on β-CD still kept the perfect intra-molecular hydrogen bonding network. The poor solubility of β-CD is believed to be attributed to this stable hydrogen bonding network and methylation is used to break this network to increase the solubility [39]. However, the hydroxyl groups on dimethyl-β-CD are not able to interact with the carbonyl group on 2a either. Because of lack of favorable hydrogen bonding interactions, the overall electrostatic contribution (ΔE_Coulomb_ + ΔE_PB_) is positive for the binding between the reactants and cyclodextrins. In sharp contrast to the hypothesis that Hbonds between the carbonyl groups of 2a and cyclodextrins play the key role in the catalysis reaction between compound 1 and maleimide derivatives including 2a [15], VM2 results do not support this hypothesis.

**Figure 4.**
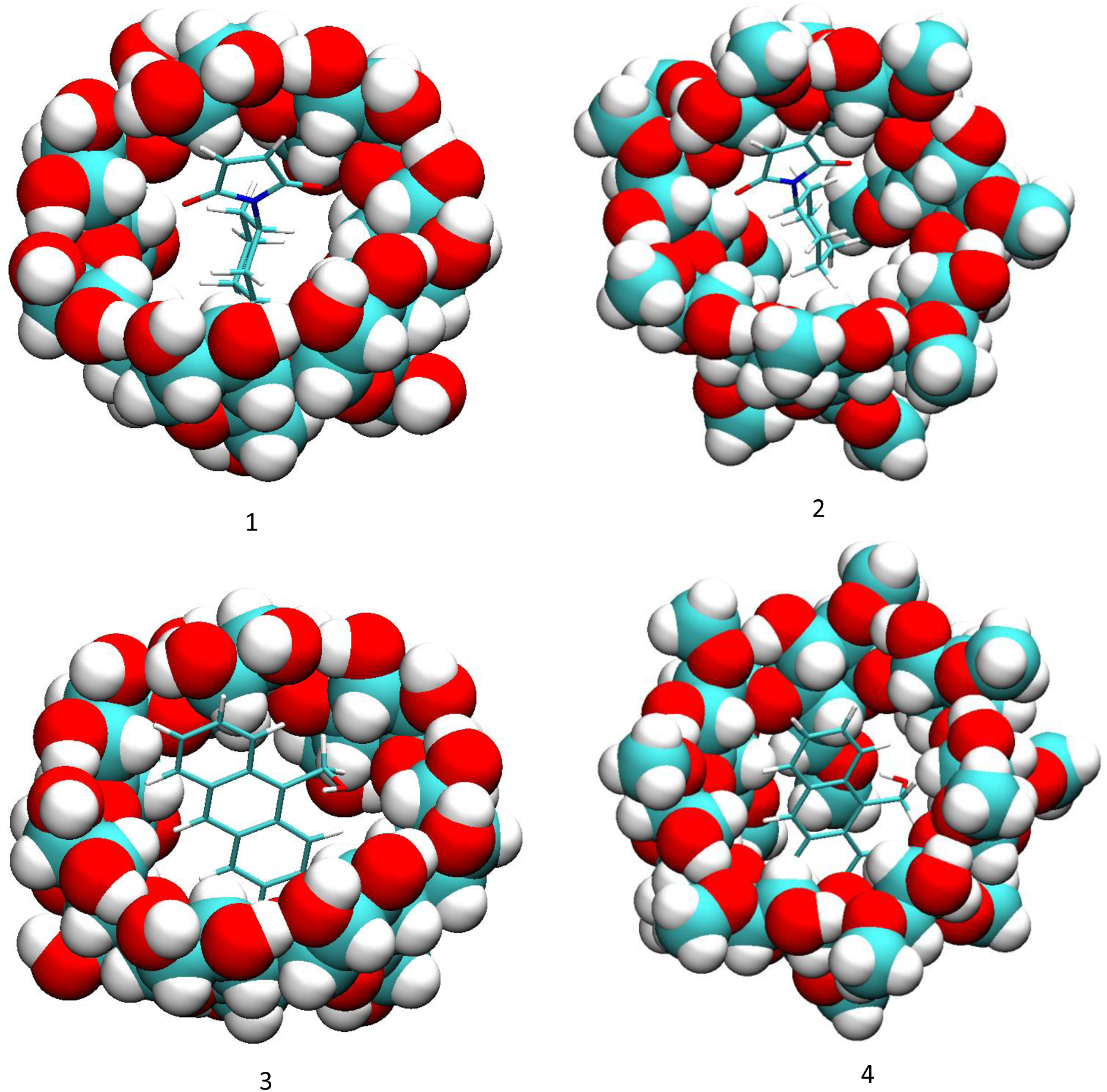
Binding modes of compounds 1 and 2a to cyclodextrins by VM2. (1) 2a and β-CD; (2) 2a and dimethyl-β-CD; (3) 1 and β-CD; (4) 1 and dimethyl-β-CD.

### 3.2 Conformational fluctuations modeled by MD simulations

The VM2 method provides multiple local and global energy minima conformations, which also reveals that the systems are highly flexible. In solution, the systems rarely stay in their energy minima conformation, and each molecule constantly fluctuates. Therefore, we ran 300-ns all-atom unbiased MD simulations in an explicit solvent model for the complexes and found more diverse conformations.

We first examined the binding strengths and conformations of the complexes of 2a and β-CD–dimethyl-β-CD and the results confirmed the stronger enthalpic interaction between 2a and dimethyl-β-CD than with β-CD due to more favorable van der Waals interactions. We calculated the interaction energies between the reactant and cyclodextrins with the MMPB/SA method to examine their intermolecular attractions (Table 2). The MMPB/SA energies are consistent with the VM2 results in that dimethyl-β-CD has stronger enthalpy contribution to the binding with 2a than β-CD. The dominant contributor is the van der Waals interaction, and the overall electrostatic interaction is positive. Table 3 compares the average representative distances between the geometric center of 2a or its moieties and the center of a cyclodextrin. Larger positive distances indicate that a reactant locates farther from the center of the cyclodextrin cavity and near the wide rim of a cyclodextrin. In contrast, a small or negative value indicates that the reactant 2a binds deeply into the pocket of a cyclodextrin toward the narrow rim. In general, 2a binds deeper into non-polar pocket of β-CD than that of dimethyl-β-CD sampled by both methods. The MD sampling provides reasonable binding modes for 2a, and the cyclohexyl group of 2a forms a close contact inside the non-polar pocket of cyclodextrins. In contrast with 2a binding to β-CD, the compound binds near the top of dimethyl-β-CD, and the shallow binding pose allows the 5-membered maleimide ring to expose more to the solvent, thereby increasing the possibility for compound 1 to interact with 2a and dimethyl-β-CD. This binding mode also brings the carbonyl group to a closer proximity to the secondary hydroxyl groups of dimethyl-β-CD to possibly form an Hbond.

**Table 2.** The MMPB/SA energies and breakdowns for the complexes of reactants and cyclodextrins. Unit in kcal/mol.

| | $\Delta E_{\text{Coulomb}}$ | $\Delta E_{\text{PB}}$ | $\Delta E_{\text{NP}}$ | $\Delta E_{\text{VDW}}$ | $\Delta H_{\text{MMPB/SA}}$ |
| --- | --- | --- | --- | --- | --- |
| β-CD with 2a | -7.24 | 17.23 | -2.19 | -23.07 | -15.27 |
| dimethyl-β-CD with 2a | -3.38 | 13.35 | -2.18 | -24.51 | -16.72 |
| β-CD with 1 and 2a | -16.00 | 27.79 | -2.66 | -25.88 | -16.75 |
| dimethyl-β-CD with 1 and 2a | -5.51 | 19.17 | -2.96 | -31.09 | -20.39 |

**Table 3.** Binding modes and conformation fluctuations sampled by MD and VM2. The average distances between geometric centers of 2a or its moieties and the geometric center of cyclodextrins in the course of 300 ns MD runs and the VM2 results. Unit in Å. In the illustration above the table, black dot represents the center of cyclodextrins, and red dot represents the center of 2a or its moieties; d1 is the distance between the centers of 2a and cyclodextrins; d2 is the distance between the centers of cyclohexyl group and cyclodextrins; d3 is the distance between the centers of imide moiety and cyclodextrins. Larger positive distances indicate that the ligand locates farther from the center of the cyclodextrin cavity.

|  | MD |  | VM2 |  |
| --- | --- | --- | --- | --- |
| β-CD | d1 | $1.44 \pm 0.70$ | d1 | $0.38 \pm 0.11$ |
| | d2 | $-0.03 \pm 0.87$ | d2 | $-1.39 \pm 0.15$ |
| | d3 | $3.35 \pm 0.68$ | d3 | $2.14 \pm 0.11$ |
| dimethyl-β-CD | d1 | $2.05 \pm 0.73$ | d1 | $1.42 \pm 0.13$ |
| | d2 | $0.67 \pm 0.87$ | d2 | $-0.10 \pm 0.17$ |
| | d3 | $3.95 \pm 0.70$ | d3 | $3.37 \pm 0.12$ |

We examined the proportion of Hbonds between N-cyclohexylmaleimide and the hydroxyl groups of cyclodextrins during the MD simulations and found very few. This observation does not support the hypothesis that these Hbonds play an important role in the catalysis as claimed by Chaudhuri *et al* [15]. For the dimethyl-β-CD and 2a complex, the proportion is negligibly 0.06% and for β-CD, it is 3.23%. The very low percentage indicates that these Hbonds are not stable, which is consistent with VM2 results, with no such Hbonds shown in all energy minima. Our results suggest a different mechanism other than a substrate forming the Hbonds with cyclodextrins to promote the Diels-Alder reaction. Assuming that the Hbonds play an important role in enhanced catalysis, our results show that β-CD can have a slightly higher possibility of forming Hbonds, which should result in a better catalyst. However, experiments did not support this hypothesis, either.

We then investigated the simultaneous binding of compounds 1 and 2a to β-CD or dimethyl-β-CD. The MMPB/SA results show that the binding of 1 and 2a with cyclodextrins has more favorable energy than the binding of 2a alone (Table 2), so after 2a binds to the cavity of cyclodextrins, 1 would also spontaneously bind to the complex. Therefore, cyclodextrins can assist in geometrically pre-organizing the two reactants for the Diels-Alder reaction. The binding is driven by van der Waals interaction. Dimethyl-β-CD binds 1 and 2a more tightly than does β-CD because it has more non-polar groups to contribute van der Waals energy. Figure 5 shows the typical pose of 1 when interacting with 2a and dimethyl-β-CD. One end of its anthracene ring and the maleimide ring of 2a forms π-π stacking, and the other end contacts with the methyl-oxyl group on dimethyl-β-CD. We further inspected the dynamics of compound 1 in the complexes. Binding of 1 to the complex of 2a and CDs is not as stable as binding to the CDs without 2a because 1 frequently moved in and out of the binding site in the complex of 2a and CDs. 2a and 1 are more stable in dimethyl-β-CD than β-CD. The tri-molecule complex with dimethyl-β-CD was present at 34.9% of the MD simulation time, which is significantly longer than the tri-molecular complex with β-CD, which is only present at 8.1% of the MD time (Figure SI 2). In addition to the percentage, we examined the length of formation of the tri-molecular complex. The 1–2a– dimethyl-β-CD complex can last as long as 60 ns, in contrast with the 1–2a–β-CD complex which formed for less than 10 ns. Figure 6 records the locations of the geometric center of 1 with the initial conformation of 2a and cyclodextrins as a reference during 300 ns. Notably, the initial conformations are the bound states of both reactants and β-CD/dimethyl-β-CD, as described in Section 2.2. These results suggest that dimethyl-β-CD is able to keep both reactants in its binding site for much longer than β-CDs.

**Figure 5.**
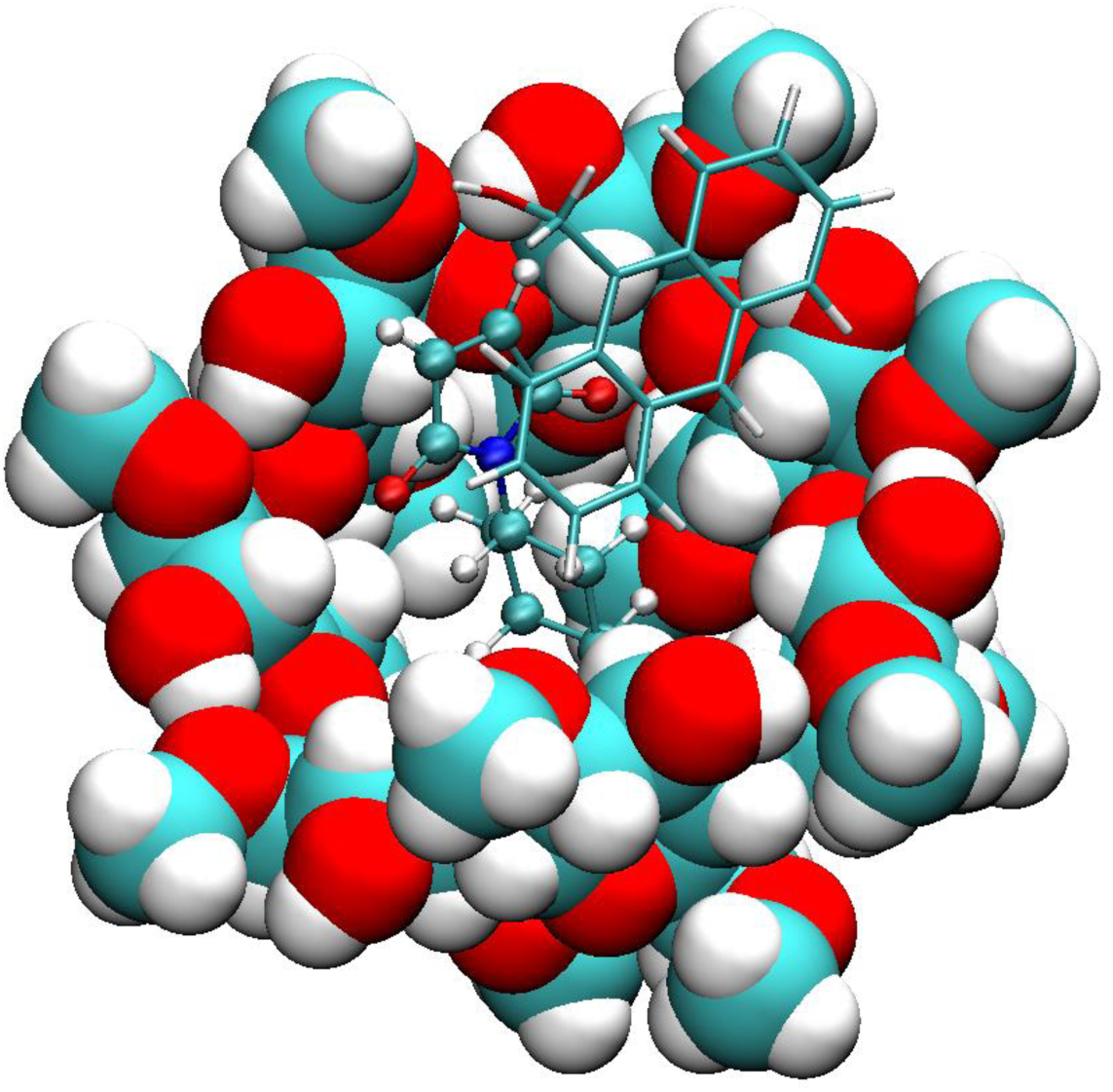
Binding mode of compound 1 with 2a and dimethyl-β-CD. Compound 1 is rendered in licorice, 2a in CPK and dimethyl-β-CD in VDW.

**Figure 6.**
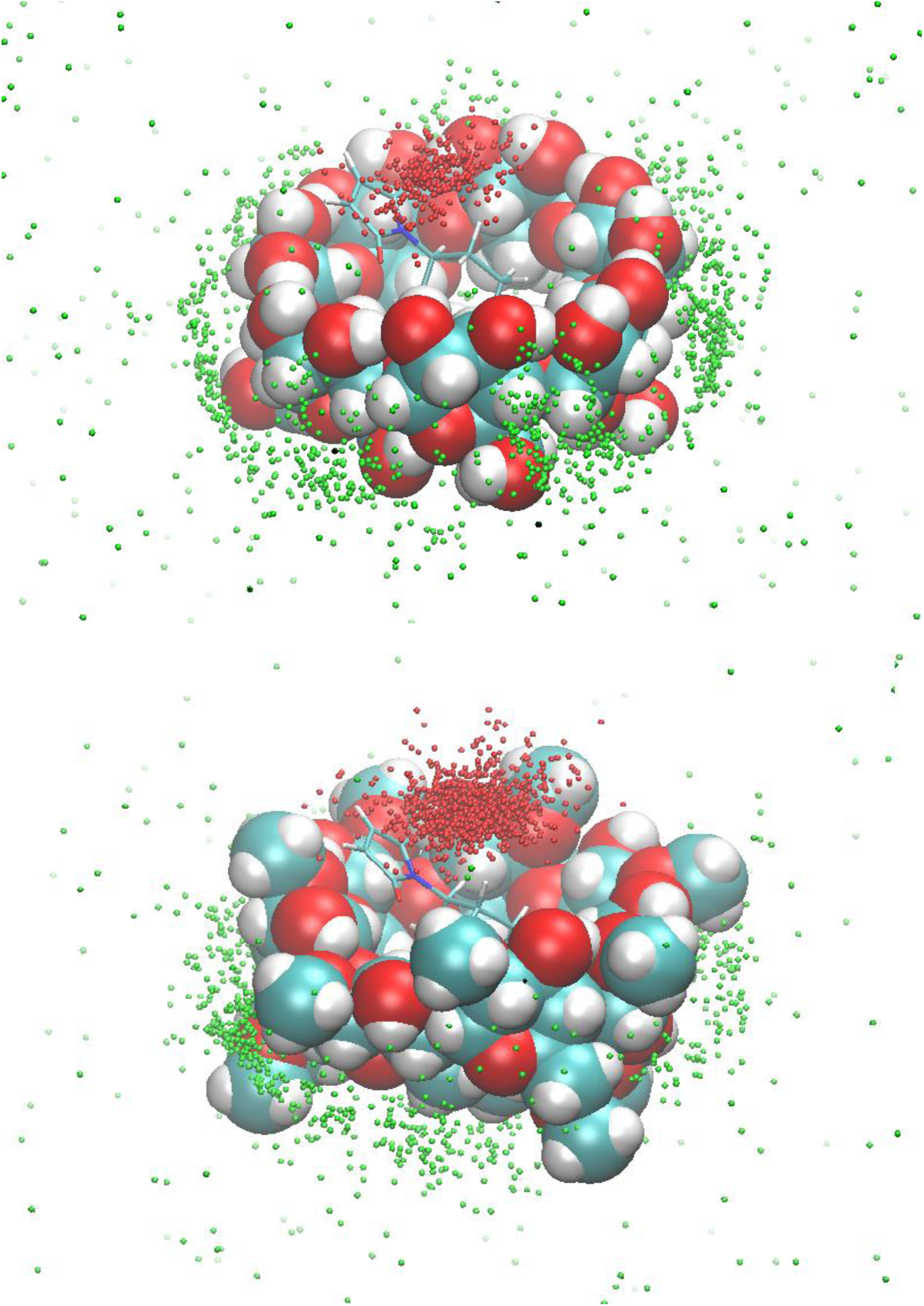
The chances of compound 1 inside the binding site of β-CD (top) and dimethyl-β-CD (bottom) in the presence of compound 2a. The red dots (inside the binding site) and the green dots (outside the binding site) are the geometric centers of compound 1 in the course of 300 ns MD runs. The initial conformations of compound 2a and β-CD/dimethyl-β-CD for the MD runs are used as reference.

### 3.3 Explanation of enhanced catalysis by QM calculations

Our molecular mechanics calculations suggest that cyclodextrins help pre-organize the two reactants in the correct geometry arrangement. To gain a better understanding of whether such a pre-organization can actually lower the activation barrier and promote a Diels-Alder reaction, we used QM calculations for five reactions: Diels-Alder reaction of 1 with 2a without cyclodextrins, and 1 and 2 plus dimethyl-β-CD or β-CD with or without the Hbond formation (Figure 7 and SI-3). As an initial effort, a semi-empirical PM3 method is used to locate the TSs and the corresponding IRCs with and without β-CD or dimethyl-β-CD. Despite its limitations, the PM3 method correctly characterizes the bond breaking and formation of Diels-Alder reactions [40] and is a practical method for large systems as in this work. Indeed, as shown in the discussion below, the reaction energetics and catalytic efficiency ranking of the catalysts predicted by PM3 agree with the experimental results.

**Figure 7.**
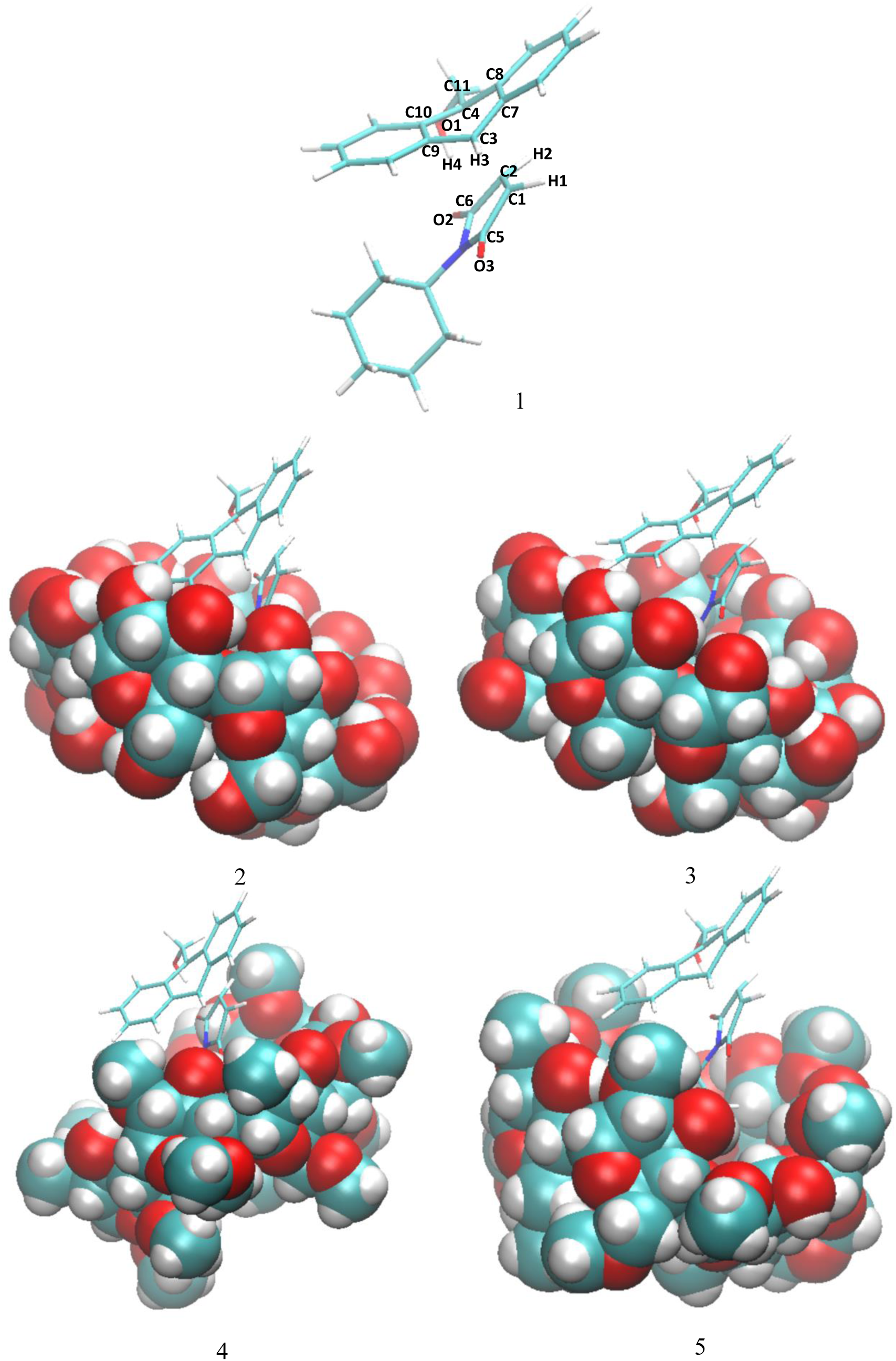
The transition states of five reactions. (1) Transition state structure in the non-catalyzed reaction between compounds 1 and 2a. Atoms names used in the table are marked on this structure. (2) Transition state structure in the reaction catalyzed by β-CD, with hydrogen bonds between the two carbonyl groups of 2a and β-CD. (3) Transition state structure in the reaction catalyzed by β-CD, without the hydrogen bonds. (4) Transition state structure in the reaction catalyzed by dimethyl-β-CD, with hydrogen bonds between the two carbonyl groups of 2a and dimethyl-β-CD.

The QM calculations suggest that Hbonds play a nearly negligible role in lowering the activation energy (ΔG in Table 4, set 1 and Figure SI 3). The presence of a cyclodextrin significantly reduced the entropy cost during the activation process, which is the main contributor to enhancing the catalysis (Table 4). Structures with the lowest energies during the MD runs were optimized with the GEDIIS method and used for searching the TS of a cyclodextrin-catalyzed reaction. The optimized structures of TSs and their key geometric parameters are depicted in Figure 7 and Table 5, and the corresponding IRCs are shown in Figure SI-3. Figure 7 shows that the non-catalyzed TS configuration is largely maintained in the catalyzed reactions. In all of the TSs, the lengths of the two new “covalent bonds” between 1 and 2a are similar, as are the improper dihedral angles at the four carbon atoms where these two new bonds are to be formed. Therefore, the catalytic effects of cyclodextrins are not from the direct participation of bond formation at TS. In addition, the IRCs with and without Hbonds with both catalysts (Figure SI-3, top and middle respectively) are similar. The reactions have slightly lower activation energies by no more than 3.0 kcal/mol when Hbonds exist between catalysts and 2a. This finding suggests that hydrogen bonding is not a significant role player in catalyzing the reaction. Surprisingly, when the IRC of the uncatalyzed reaction is plotted against that of catalyzed reactions without Hbonds (Figure SI-3 bottom), they look almost identical. The activation energies are 28.20, 28.75 and 27.28 kcal/mol for reactions without a catalyst, with β-CD, and with dimethyl-β-CD, respectively. Because IRC calculations report only the potential energies for the reaction systems, the high similarity in potential energies suggests that the catalytic power of cyclodextrins may be related to entropy.

**Table 4.** The activation free energies and its enthalpy and entropy components for the Diels-Alder reactions with and without cyclodextrin catalysts. Unit in kcal/mol.

| | $\Delta G$ | $\Delta H$ | $-T\Delta S$ |
| --- | --- | --- | --- |
| non-catalyzed reaction | 49.04 | 31.76 | 17.28 |
| reaction catalyzed by β-CD, set 1<sup>a</sup> | 36.47 | 30.96 | 5.50 |
| reaction catalyzed by β-CD, set 2<sup>b</sup> | 38.32 | 33.32 | 5.00 |
| reaction catalyzed by dimethyl-β-CD, set 1<sup>a</sup> | 29.46 | 27.90 | 1.56 |
| reaction catalyzed by dimethyl-β-CD, set 2<sup>b</sup> | 29.17 | 28.13 | 1.04 |

<sup>a</sup> Transition state structure has hydrogen bonds between the two carbonyl groups of 2a and cyclodextrins.
<sup>b</sup> Transition state structure does not have hydrogen bonds.

**Table 5.** The key geometric parameters of the transition states. Bond stands for bond length, improper for improper dihedral. Atom names are indicated in Figure 7. Unit is Å for bond length, and degree for improper dihedral angles.

|  | bond C1-C3 | bond C2-C4 | improper<br>C1-C2-H1-C5 | improper<br>C2-H2-C1-C6 | improper<br>C3-H3-C7-C9 | improper<br>C4-C8-C11-C10 |
| --- | --- | --- | --- | --- | --- | --- |
| 1 | 2.130 | 2.298 | 23.119 | 20.047 | 16.033 | 13.738 |
| 2 | 2.125 | 2.316 | 22.883 | 19.641 | 15.879 | 13.229 |
| 3 | 2.127 | 2.312 | 23.413 | 19.531 | 16.184 | 13.394 |
| 4 | 2.141 | 2.297 | 22.869 | 21.064 | 15.451 | 13.417 |
| 5 | 2.140 | 2.283 | 22.885 | 20.346 | 15.911 | 13.947 |

Thus, we calculated the activation free energy and its enthalpy and entropy components for all five reactions (Table 4). The activation free energy for the non-catalytic reaction was 49.04 kcal/mol, which is in the range of experimental activation free energies for typical Diels-Alder reactions [41, 42]. Also, in Table 4, cyclodextrins lower the energy barriers significantly and dimethyl-β-CD performs better than β-CD, in agreement with the experimental observation [15]. The calculated activation free energies can be decomposed into enthalpy and entropy terms. As shown in Table 4, the first sets of both β-CD and dimethyl-β-CD complexes have a lower enthalpy term than the second set, so the Hbonds may help stabilize the transition state. However, the presence of Hbonds lowers the free energy by only less than 2 kcal/mol for both catalysts. As compared with the 10- to 20-kcal/mol stabilization of the activation energy in the catalyzed reactions, the Hbond impact is small. Regardless of the presence of Hbonds, the TSs in the dimethyl-β-CD catalyzed reaction have lower enthalpy than those in the β-CD catalyzed reaction. Most importantly, the entropy term-TΔS is the dominant factor in differentiating the activation energies. It drops from 17.3 kcal/mol in the non-catalyzed reaction to 5.0 ∼ 5.5 kcal/mol in the β-CD catalyzed reaction, and to 1.0 ∼ 1.5 kcal/mol in the dimethyl-β-CD catalyzed reaction. Therefore, the binding of compounds 1 and 2a to cyclodextrins acts as a pre-organization and remarkably reduces the entropy penalty to the activation free energy for the transition state of 1 and 2a.

The above results show that instead of directly participating in chemical bonding at the transition state, the pre-organization provided by complexation between compounds 1 and 2a and CDs lowers the activation energy of the Diels-Alder reaction. The catalytic effect is achieved by the reduction of ΔS when going from the reactant state to the reaction barrier. The limitations of this QM study also deserve comments. Because of the existence of multiple conformers of cyclodextrins, there may be multiple reaction pathways and TS structures. Because cyclodextrins do not participate in the chemical bonding at the barrier, its configurations are not expected to change significantly during the reaction. Therefore, the configuration effect is largely cancelled in activation energy calculations. Furthermore, the initial configurations used in the QM calculations are those with the lowest energies from MD runs, so they may represent the configuration with the largest statistical weight. Nevertheless, we could further investigate the configurational contributions of cyclodextrin to the activation barrier. Solvent effects were not considered in this QM study. They are expected to be similar for both catalyzed and un-catalyzed reactions, because the reaction coordinates of the TS (i.e., eigenvectors) are similar for both catalyzed and un-catalyzed reactions. Omitting solvent effects does not affect the conclusions of this paper. In addition, PM3 is a low-level theory. Its use usually needs to be benchmarked by high-level theories such as DFT. However, our PM3 activation energies for the reaction barrier were in the range of the experiment and correctly ranked the CD catalysts. These observations suggest that the PM3 results are reasonable.

In summary, the major findings from this QM study are as follows: (1) entropy plays a significant role in lowering the activation free energy barrier in the Diels-Alder reactions catalyzed by cyclodextrins; (2) hydrogen bonding, although lowering the barrier to some extent, is less important than entropy contribution; and (3) dimethyl-β-CD is more efficient than β-CD in catalyzing the reaction.

## 4. Conclusion

Dimethyl-β-CD could catalyze Diels-Alder reactions of 9-anthracenemethanol with a variety of N-substituted maleimides under mild reaction conditions. To understand the mechanism, we performed MM and QM calculations. The driving force of the catalysis is not the Hbonds between cyclodextrins and 2a because of the low occurrence of the Hbonds suggested by VM2 and MD results, and IRC calculations yielded small difference with and without the Hbonds. However, cyclodextrins, especially dimethyl-β-CD, do show strong binding affinity with a pre-organized 1– 2a reactant complex and increase the probability of the Diels-Alder reaction. Moreover, QM calculations showed that cyclodextrins remarkably reduced the activation entropy. These findings depict a possible two-step catalysis mechanism by cyclodextrins: the first step is the complex formation between cyclodextrins and reactants and the second step is the reaction of reactants inside the cavity of cyclodextrins to form the product. The formation of the complex is the key to the catalysis because it not only increases the rate that reactant molecules would meet each other but also lowers the entropy and thus the activation free energy of this reaction. Dimethyl-β-CD outperforms β-CD because of stronger van der Waals interaction with the pre-organized reactants and better performance in reducing the activation energy. PM3 results are in qualitative agreement with experiments on activation free energies of the Diels-Alder reactions and the relative catalytic activities for the systems under study. Further validating PM3 in predicting the cyclodextrin-catalyzed Diels-Alder reactions at the quantitative level and understanding details of the catalytic mechanisms would require higher-level *ab initio* calculations. Work in this direction is being conducted.

## Acknowledgements

We thank support from the US National Institute of Health (GM-109045), US National Science Foundation (MCB-1350401), and NSF national super computer centers (TG-CHE130009). We also thank Dr. Mindy Levine for insightful discussions on using β-CDs as a catalyst for promoting chemical reactions.

## Supplement materials

**Figure SI-1.**
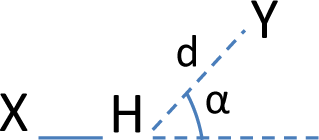
Definition of hydrogen bonds. X and Y stand for the donor and the acceptor respectively. d is the distance between the acceptor Y and the hydrogen, and α is the complimentary angle of X-H Y. A hydrogen bond is formed if d is smaller than 2.0 Å and α is smaller than 90°.

**Figure SI-2.**
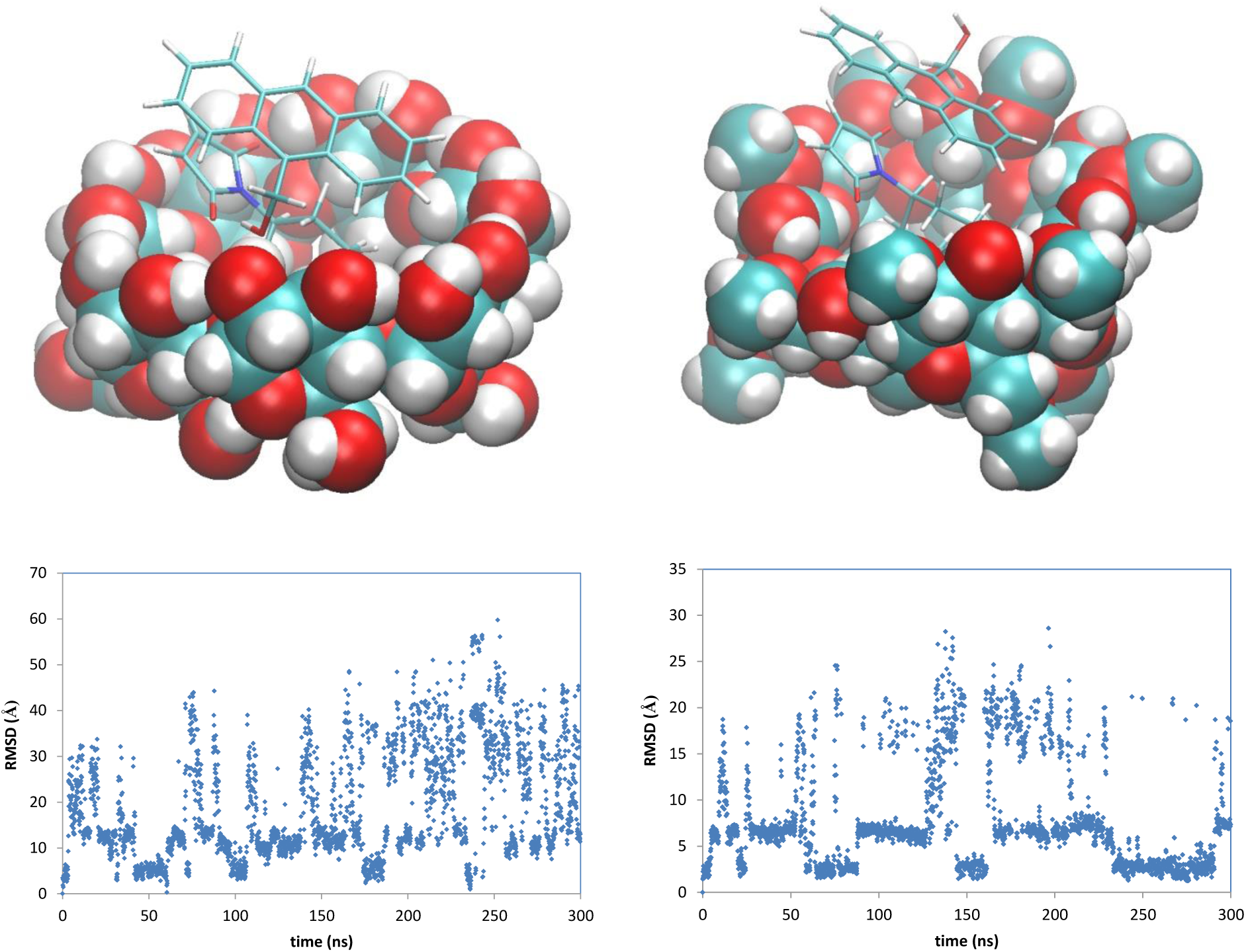
RMSD (bottom) of compound 1 with respect to the initial conformations (top) in the 300 ns MD runs. Left: complex of β-CD with 1 and 2a; right: complex of dimethyl-β-CD with 1 and 2a.

**Figure SI-3.**
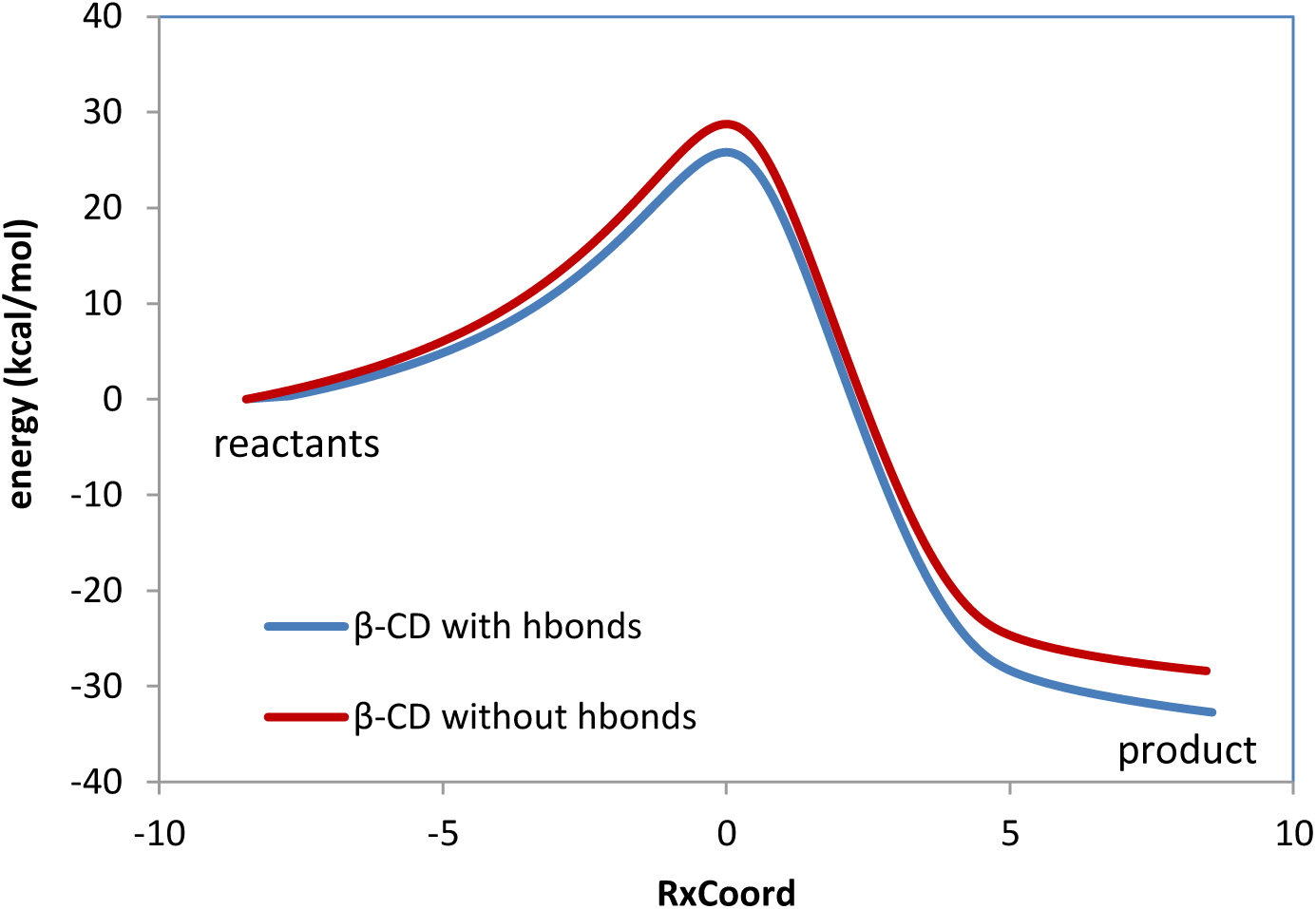
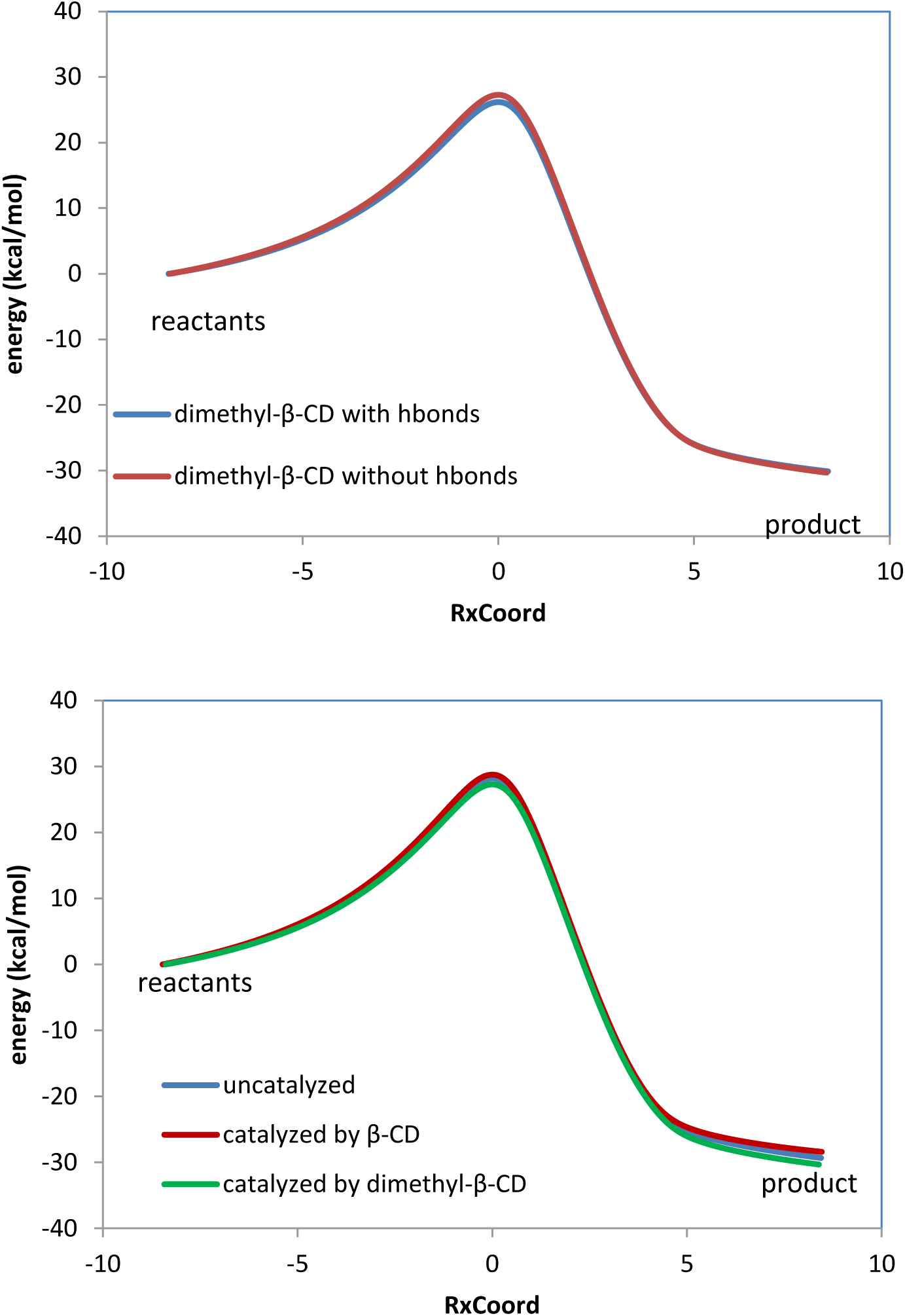
Reaction paths obtained by IRC. In all plots the curves are normalized to set the reactants energies at zero point. Top: reaction paths for the reaction catalyzed by β-CD, with and without the hydrogen bonds. Middle: reaction paths for the reaction catalyzed by dimethyl-β-CD, with and without the hydrogen bonds. Bottom: reaction paths for uncatalyzed reaction between compounds 1 and 2a, reaction catalyzed by β-CD and reaction catalyzed by dimethyl-β-CD. No hydrogen bonds exist between compound 2a and cyclodextrins.

